# Fast Low-Input Efficient Hi-C

**DOI:** 10.1101/2024.08.28.609956

**Authors:** Mor Guetta, Nili Avidan, Hagai Kariti, Liron Davis, Revital Shemer, Evan Elliot, Noam Kaplan

## Abstract

We developed FLIE (Fast Low-Input Efficient) Hi-C, a highly optimized and simplified version of Hi-C. Our method shortens the duration of Hi-C from five days to two days and decreases key reagent amounts 5-20 fold, while maintaining experimental controls and data quality. Importantly, the method also reduces the required input material by 100-1000 fold, as we demonstrate by generating detailed high complexity interaction maps from low-input sorted mouse frozen neuron nuclei and from 5,000-10,000 human Hap1 cells.

## Main

Hi-C is a powerful genomic technique for studying spatial genome organization, based on measuring spatial interactions of pairs of genomic loci in high throughput using next generation sequencing^1–3^. In the past decade, the popularity of Hi-C has risen due to the increasing appreciation of the role of genome organization in genome function^4–6^, but also due to the discovery that Hi-C can be used to boost genome assembly^7–10^. Together, these notions suggest potentially leveraging Hi-C to gain an integrated genetic (e.g. structural variation) and epigenetic mapping of clinical samples^11–13^. Despite its utility in diverse applications and despite important improvements introduced over the years^14–17^, Hi-C remains a technically challenging and costly endeavor, especially when starting material is limited.

Following the initial Hi-C experiments, two major improvements introduced in the second generation of Hi-C were changing the restriction enzyme from HindIII (a 6 cutter) to DpnII/MboI (a 4 cutter) and performing the key experimental steps in-nucleus, together enhancing the resolution and quality of maps^2,3^. This version of Hi-C has been used widely, including in the 4D Nucleome and ENCODE consortiums^18,19^.

Here we introduce a highly optimized Hi-C version called FLIE (Fast Low Input Efficient) Hi-C, based on the Dekker lab’s Hi-C 2.0 protocol^3^. As in Hi-C 2.0, we use DpnII as the main restriction enzyme and provide quality controls at stages in the experiment (chromatin integrity, DpnII digestion, ligation, biotin pulldown, library preparation and ClaI digestion) allowing to pinpoint technical issues. We introduced several improvements including the omission and reordering of steps, reduction of incubation times and reduction of volumes and reagents. Together, these lead to a markedly shorter protocol which can comfortably been completed in two full days, loses less material, and offers a ∼5-20 fold reduction in amounts of key reagents while producing high-quality interaction maps.

We briefly overview the protocol and its main improvements with respect to Hi-C 2.0 (**Table 1**). Throughout the experiment, we greatly reduce reaction volumes and eliminate several washing and filtering steps, leading to a ∼5-20 fold decrease in the amounts of key reagents (**Table 2**). The first day of FLIE Hi-C starts with formaldehyde crosslinking of cells, followed by cell lysis. We skip dounce homogenization used in Hi-C 2.0, and proceed to DpnII digestion for 1.5 hours (rather than overnight). We then fill in digested ends with biotinylated dATP and blunt end ligation, both performed 1 hour each (instead of 4 hours each). Next, crosslinks are reversed for 1 hour (rather than 2 hours+overnight). DNA is then purified and quantified by a simplified protocol, followed by biotin removal from unligated DNA ends performed for 1 hour (rather than 4) and left overnight, completing the first day in ∼8 hours. The second day of FLIE Hi-C starts with DNA sonication. To minimize loss of material, FLIE Hi-C skips size selection and end-repair and progresses directly to biotin pulldown by streptavidin beads. Instead of separate end-repair, A-tailing and adapter ligation, we use the NEBNext Ultra II library preparation kit which combines these steps. At this stage, the sample is ready for calibration PCR to identify the minimal number of PCR cycles necessary to produce the Hi-C library. We use this amplified sample to also perform quality control by ClaI digestion (rather than at the end as in Hi-C 2.0). If the library is properly digested by ClaI, we proceed to final production PCR amplification of the library, size selection and cleanup. The second day consists of approximately 5.5 hours.

**Table 1.** Approximate timeline for FLIE Hi-C versus Hi-C 2.0. * -End repair precedes biotin pulldown in Hi-C 2.0. ** -End repair, A tailing and adapter ligation are performed simultaneously in FLIE Hi-C. *** -Production PCR precedes ClaI quality control in Hi-C 2.0.

| Experimental stage | FLIE Hi-C | Hi-C 2.0 |
| --- | --- | --- |
| **Crosslinking** | 1 hour [day 1] | 1 hour |
| **Lysis and digestion** | 3 hours [day 1] | 1 hour + overnight |
| **Biotin fill-in** | 1 hour [day 1] | 5 hours |
| **Ligation** | 1 hour [day 1] | 4 hours |
| **Reverse crosslinking** | 1 hour [day 1] | 2 hours + overnight |
| **DNA purification** | 1 hour [day 1] | 2.5 hours |
| **ClaI+MboI quality control** | skipped | 2 hours |
| **Biotin removal** | overnight | 4.5 hours |
| **Sonication** | 0.5 hour [day 2] | 0.5 hour |
| **Size selection** | skipped | 1 hour |
| **Biotin pulldown\*** | 0.5 hour [day 2] | 0.5 hour |
| **End repair\*** | ** | 1 hour |
| **A tailing** | 1.5 hour [day 2] | 1 hour |
| **Adapter ligation** | ** | 2.5 hours |
| **Calibration PCR** | 1 hour [day 2] | 1 hour |
| **ClaI quality control\*\*\*** | 1 hour [day 2] | 1 hour |
| **Production PCR\*\*\*** | 1 hour [day 2] | 1 hour |
| **Total time** | **13.5 + 1 night** | **31.5 hours + 2 nights** |

**Table 2.** Reduction in amounts of key reagents in FLIE Hi-C versus Hi-C 2.0. All reagents are identical between FLIE Hi-C and Hi-C 2.0 except for T4 DNA ligase, where we verified that ligation activity per µl is approximately equal.

| Step | Reagent | FLIE Hi-C | Hi-C 2.0 | Fold reduction |
| --- | --- | --- | --- | --- |
| **Chromatin digestion** | DpnII | 50U | 400U | 8 |
| **Biotin fill-in** | Polymerase I<br>Klenow | 10U | 50U | 5 |
| **Biotin fill-in** | Biotinylated dATP | 2 nmol | 15 nmol | 7.5 |
| **Ligation** | T4 DNA ligase | 2.5 µl | 50 µl | 20 |
| **Biotin removal** | T4 DNA<br>polymerase | 9U | ~45U | ~5 |

To test FLIE Hi-C in a realistic limited-input setting with primary biological material, we applied the method to frozen sorted mouse neuron nuclei, altogether four samples of 50,000 nuclei each. After verifying the experimental quality controls, we sequenced, mapped and processed the data to generate interaction maps (see Methods). The interaction map metrics indicated excellent data quality^20^, with 86% cis/trans interaction ratio and <3% PCR duplicates, suggesting that the libraries are high quality and that at this level of sequencing library complexity is not a limiting factor. To study the interaction maps visually, we pooled the four samples, resulting in an interaction map generated from a total of 128M unique mapped read pairs and compared to a map generated from one sample with only 35M unique mapped read pairs. Upon visual examination, we find that despite the modest sequencing depth, small scale structures such as TADs and point interactions are easily discernible at high resolutions (**Figure 1a**).

**Figure 1.**
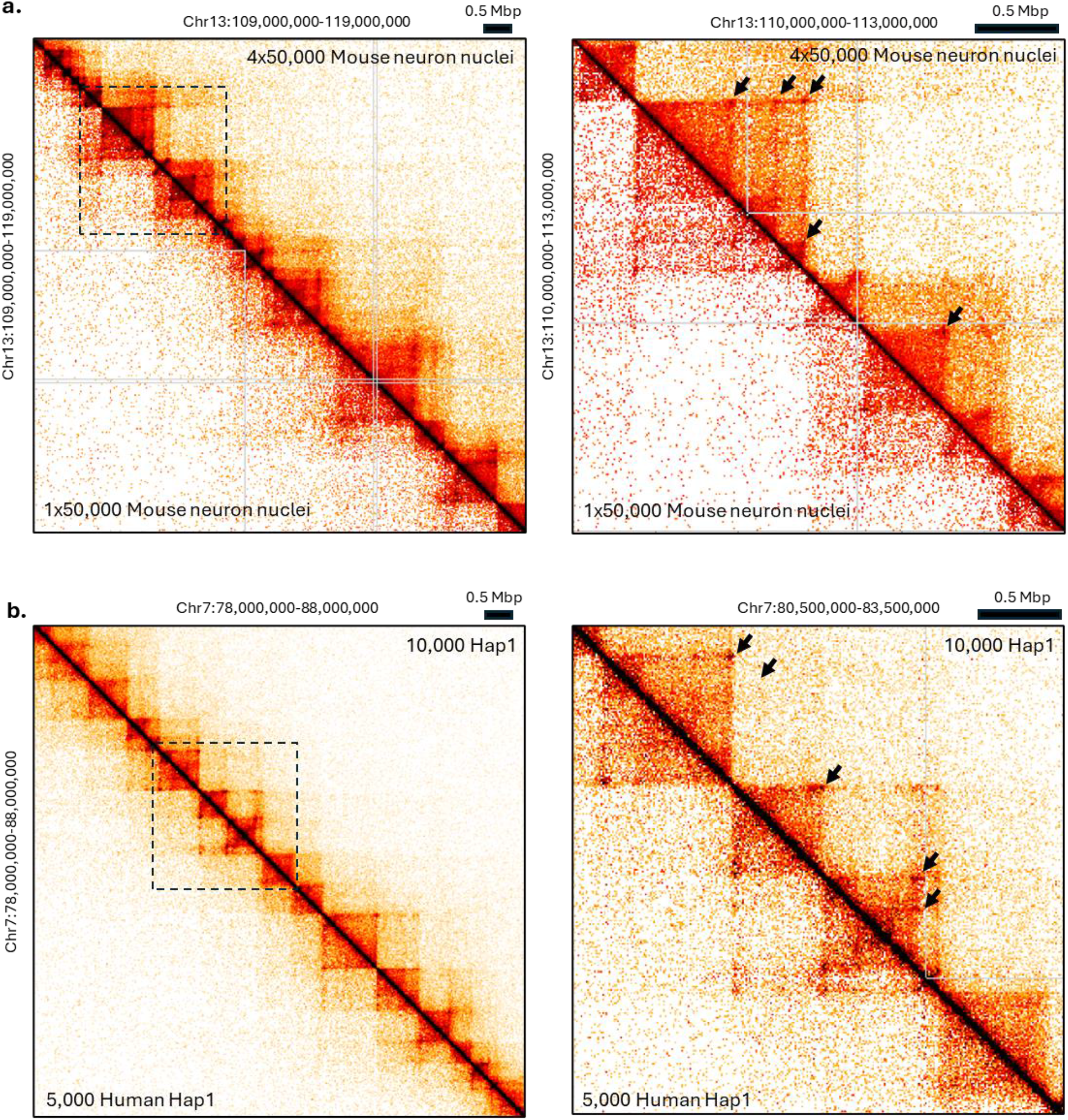
FLIE Hi-C interaction maps from moderately sequenced low-input samples. **a.** Interaction maps of mouse neuron nuclei (1×50,000 vs. 4×50000 nuclei with 35M and 128M unique mapped reads respectively). Left: 10 Mbp genomic region at 25 Kbp resolution showing detailed TAD structure. Right: Zoomed in 3 Mbp region at 10 Kbp resolution showing point interactions (black arrows). **b**. Detailed interaction maps generated from human haploid Hap1 cells with (5,000 vs 10,000 cells with 124M and 195M unique mapped reads respectively). Left: 10 Mbp genomic region at 25 Kbp resolution showing detailed TAD structure. Right: Zoomed in 3 Mbp region at 10 Kbp resolution showing point interactions (black arrows).

Finally, we investigated if FLIE Hi-C could produce interaction maps from ultra-low cell amounts. To this end, we applied FLIE Hi-C to 5,000 and 10,000 FACS-sorted human haploid Hap1 cells. This represents a 100-1000 fold reduction relative to typical second generation Hi-C requirements. We successfully produced libraries from these samples, which passed quality control. The overall quality of the data was high, with >88% cis. As expected, library complexity was lower than in the mouse neuron nuclei experiments, with 14% estimated PCR duplicates in the 10,000 cell sample and 16% estimated PCR duplicates in the 5,000 cell sample (after removing optical/machine duplicates, see Methods). Still, this complexity level can support significantly deeper sequencing of the samples. Examining the interaction maps visually, we find that at this sequencing depth (124M and 195M mapped unique read pairs), TAD structure and some point interactions are evident (**Figure 1b**).

We conclude that FLIE Hi-C constitutes a significant enhancement of second generation Hi-C, requiring less time, reagents and input material. This simple and efficient Hi-C protocol serves as an excellent platform for incorporating further variations proposed by others (e.g. alternative crosslinking and digestion approaches^16,17^), for developing derivative methods, for studying clinical or other biologically-precious samples, and for performing Hi-C in parallel at large scale.

## Methods

### Sample preparation

Mouse forebrain was dissected and minced using homogenizer in 5 ml homogenization buffer (0.25M sucrose, 25mM KCl, 5mM MgCl2, 20mM Tricine-KOH, 1mM DTT, 0.15mM spermine, 0.5mM spermidine), 7 µl RNase Inhibitor (Promega N2611) and 300 µl 5% IGEPAL. The sample was filtered through a 40 µm strainer, mixed with 5 ml of 50% iodixanol (Sigma D1556), underlaid with a gradient of 30% and 40% iodixanol, and centrifuged at 4,600g for 1 hour in a swinging bucket centrifuge at 4°C. Nuclei were collected at the 30%-40% interface. The nuclei were transferred to a 2 ml tube and centrifuged 10 min at 1,000g and 4°C. Nuclei were then FACS-sorted into 4 tubes of 50,000 nuclei each. Finally, nuclei were transferred to freezing buffer (0.32M sucrose, 5mM CaCl2, 0.1mM EDTA, 10mM tris HCl, 3mM Mg(Ac)2) and frozen in -80°C.

5,000 and 10,000 Hap1 human haploid cells (Horizon Discovery) were FACS-sorted to ensure accurate cell counts following formaldehyde crosslinking and quenching as described in the FLIE Hi-C protocol.

### FLIE Hi-C

We provide a detailed FLIE Hi-C protocol in the Supplementary Methods. For ultra-low input (5,000 and 10,000 Hap1 cells), we adjust a few steps as detailed in the Supplementary Methods. Briefly: We avoid running chromatin integrity, digestion and ligation controls on gel; We do not quantify DNA prior to its amplification; Instead of PCR calibration and library amplification, we run 10 cycles of PCR amplification on the entire sample (rather than half), quantify the DNA and if too low (which was not the case here) amplify further, AMpure XP size-select, and quantify the final library concentration; To retain the crucial ClaI digestion control, we recycle the streptavidin beads after removing the amplified PCR product and repeat PCR on the recycled beads, and finally use the product to if the library is digested by ClaI.

FLIE Hi-C libraries from mouse neuron nuclei were sequenced on a NextSeq550 (paired-end 2×75 bp), and FLIE Hi-C libraries from Hap1 cells were sequenced on a NovaSeq6000 (paired-end 2×50 bp).

### Computational analysis

Hi-C libraries were processed using the distiller-nf pipeline (Version 0.3.4, https://github.com/open2c/distiller-nf) to generate interaction maps in mcool format^21^. Briefly, paired-end reads were aligned to the human genome (Hap1 to hg38) or the mouse genome (mouse neuron nuclei to mm10), uniquely mapping non-duplicate read pairs were retained, reads were binned into specified resolutions and interaction maps were filtered and balanced^22^. All other Hi-C analysis was implemented in custom software. We used HiGlass ^23^ to visualize and explore the data.

Due to underloading (not due to lack of library material), the NovaSeq6000 sequencing run had a high level of machine/optical duplicates. To distinguish PCR duplicates (which are needed to estimate library complexity) from machine/optical duplicates, we extracted flowcell tile information from fastq headers of duplicate reads. Given that there are 156 different tiles in the flowcell, it is highly unlikely that two reads would be equivalent (i.e. duplicates) and be located in the same tile, and we designated these as machine/optical duplicates. Using this approach we estimated that 60-70% of the duplicates were machine/optical duplicates, removed these, and corrected the read statistics accordingly.

## Data availability

Raw sequencing data and processed interaction maps (mcool) are deposited in GEO under GSEXXXXXX.

## Acknowledgements

We thank David Cohen, the Technion Biomedical Core Facility genomics and imaging teams, and past and present members of the Kaplan lab. This research was funded by an Azrieli Early Career Faculty fellowship and an Israel Science Foundation Personal Grant (1479/18) (NK).

## References

1. Lieberman-Aiden, E. et al. Comprehensive mapping of long-range interactions reveals folding principles of the human genome. Science 326, 289–93 (2009).

2. Rao, S. S. P. et al. A 3D map of the human genome at kilobase resolution reveals principles of chromatin looping. Cell 159, 1665–80 (2014).

3. Belaghzal, H., Dekker, J. & Gibcus, J. H. Hi-C 2.0: An optimized Hi-C procedure for high-resolution genome-wide mapping of chromosome conformation. Methods 123, 56–65 (2016).

4. Dekker, J. & Mirny, L. The 3D Genome as Moderator of Chromosomal Communication. Cell 164, 1110–1121 (2016).

5. Lupiáñez, D. G. et al. Disruptions of topological chromatin domains cause pathogenic rewiring of gene-enhancer interactions. Cell 161, 1012–1025 (2015).

6. Mccord, R. P., Kaplan, N. & Giorgetti, L. 3C and beyond : towards an integrative view of chromosome structure and function. Mol. Cell 1–30 (2020) doi:10.1016/j.molcel.2019.12.021.

7. Kaplan, N. & Dekker, J. High-throughput genome scaffolding from in vivo DNA interaction frequency. Nat. Biotechnol. 31, 1143–1147 (2013).

8. Burton, J. N. et al. Chromosome-scale scaffolding of de novo genome assemblies based on chromatin interactions. Nat. Biotechnol. 31, 1119–25 (2013).

9. Marie-Nelly, H. et al. High-quality genome (re)assembly using chromosomal contact data. Nat. Commun. 5, 5695 (2014).

10. Oddes, S., Zelig, A. & Kaplan, N. Three invariant Hi-C interaction patterns: Applications to genome assembly. Methods 142, 89–99 (2018).

11. Akdemir, K. C. et al. Disruption of chromatin folding domains by somatic genomic rearrangements in human cancer. Nat. Genet. 52, 294–305 (2020).

12. Dixon, J. R. et al. Integrative detection and analysis of structural variation in cancer genomes. Nat. Genet. 50, 1388–1398 (2018).

13. Xu, Z. et al. Structural variants drive context-dependent oncogene activation in cancer. Nature 612, 564–572 (2022).

14. Díaz, N. et al. Chromatin conformation analysis of primary patient tissue using a low input Hi-C method. Nat. Commun. 9, 4938 (2018).

15. Zhang, C. et al. tagHi-C Reveals 3D Chromatin Architecture Dynamics during Mouse Hematopoiesis. Cell Rep. 32, 108206 (2020).

16. Hsieh, T. H. S., Fudenberg, G., Goloborodko, A. & Rando, O. J. Micro-C XL: Assaying chromosome conformation from the nucleosome to the entire genome. Nat. Methods 13, 1009–1011 (2016).

17. Lafontaine, D. L., Yang, L., Dekker, J. & Gibcus, J. H. Hi-C 3.0: Improved Protocol for GenomeWide Chromosome Conformation Capture. Curr. Protoc. 1, 1–28 (2021).

18. Reiff, S. B. et al. The 4D Nucleome Data Portal as a resource for searching and visualizing curated nucleomics data. Nat. Commun. 13, 1–11 (2022).

19. Bernstein, B. E. et al. An integrated encyclopedia of DNA elements in the human genome. Nature 489, 57–74 (2012).

20. Lajoie, B. R., Dekker, J. & Kaplan, N. The Hitchhiker’s Guide to Hi-C Analysis: Practical guidelines. Methods 72, 65–75 (2014).

21. Abdennur, N. & Mirny, L. A. Cooler: Scalable storage for Hi-C data and other genomically labeled arrays. Bioinformatics 36, 311–316 (2020).

22. Imakaev, M. et al. Iterative correction of Hi-C data reveals hallmarks of chromosome organization. Nat. Methods 9, 999–1003 (2012).

23. Kerpedjiev, P. et al. HiGlass: Web-based visual exploration and analysis of genome interaction maps. Genome Biol. 19, 1–12 (2018).

