## Supplementary Methods for "Fast Low-Input Efficient Hi-C"

### FLIE Hi-C protocol

The protocol has been tested on samples of 5,000-3,000,000 mammalian cells.

#### 1. Cell crosslinking

1.1. Aspirate the medium, wash with 1 ml trypsin and add 1 ml of trypsin per plate to detach the cells if adherent.

1.2. Transfer cells to a 50 ml tube if adherent or just transfer cells if in suspension.

   i. Spin down cells 300 x g for 5 minutes.

   ii. Wash twice with PBS.

   iii. Count the cells and re-suspend in PBS at a concentration of 0.5-1M cells/ml.

1.3. Crosslink the cells by adding 60 µl/ml of 37% Formaldehyde to obtain 2% final concentration. Mix by gently inverting the tube a few times immediately after adding formaldehyde.

1.4. Incubate the tubes at room temperature (RT) for 10 min on a rocking platform.

1.5. Add 80 µl/ml 2.5 M Glycine to quench the crosslinking. Mix well by inverting the tube.

1.6. Incubate for 5 min at RT and then incubate on ice for 15 min to stop the crosslinking process.

1.7. Centrifuge the crosslinked samples 800 x g at 4°C for 10 min.

1.8. Wash twice with cold PBSx1 (800 x g at 4°C for 10 min).

1.9. Aliquot the cells by transferring up to 3 mil cells to 1.7 ml Eppendorf tubes.

1.10. Centrifuge the crosslinked samples 800 x g at 4°C for 10 min to remain with cell pellets.

- Cell pellets can be stored at -80°C for up to 1.5 years.

#### 2. Cell lysis and chromatin digestion with DpnII

2.1. Resuspend crosslinked cells in 20 µl cold lysis buffer and add 2 µl protease inhibitor cocktail x100.

2.2. Incubate on ice for 15 minutes.

2.3. Add 20 µl of 1% SDS and mix carefully by pipetting and avoid forming bubbles.

2.4. Incubate at 65°C for 10 minutes in a ThermoMixer (800 rpm). Spin down gently and transfer to ice for 15 min.

2.5. Add 120 µl water, 20 µl 10 x NEBbuffer3, and 20 µl 10% Triton x-100 (by this order), to a total volume of 200 µl. Mix gently by pipetting and avoid forming bubbles.

2.6. Incubate at 37°C for 15 minutes in a ThermoMixer (800 rpm).

2.7. * Skip this stage if starting material is less than 50k cells: Transfer 5 µl from the sample to a new 1.7 ml Eppendorf tube for **Chromatin Integrity control (CI)**. Keep in 4°C.

2.8. Add 50U (5 µl of 10,000 units/ml) DpnII to the Hi-C tube and mix gently.

2.9. Incubate at 37°C for 1.5 hours in a ThermoMixer (800 rpm) to digest the chromatin.

2.10. Incubate at 65°C for 20 minutes to deactivate DpnII.

2.11. * Skip this stage if starting material is less than 50k cells: Transfer 5 µl from the sample to a new 1.7 ml Eppendorf tube for **Digestion (D) control**. Keep in 4°C.

#### 3. Biotin-filling to DNA ends:

3.1. Prepare Biotin Fill-in mix:

| **Fill-in mix** | **Per sample** |
|---|---|
| 10 x NEB 3.1 buffer | 1 µl |
| 25mM dCTP & dGTP& dTTP mix | 1 µl |
| 1mM biotin-14-dATP | 2 µl |
| 5U/µl DNA polymerase I Klenow | 2 µl |
| Water | 4 µl |
| Total | 10 µl |

3.2. Add 10µl of the mix to each Hi-C tube (200 µl total). Mix gently by pippeting and avoid forming bubbles.

3.3. Incubate at 24°C for 1 hour in a ThermoMixer (800 rpm).

#### 4. Blunt end ligation

4.1. Prepare Blunt-end ligation mix:

| **Ligation mix** | **Per sample** |
|---|---|
| 10 x ligation buffer | 25 µl |
| 50% PEG-4000 | 20 µl |
| 20 mg/ml BSA | 1.25 µl |
| T4 DNA ligase 400 U/ µl | 2.5 µl |
| water | 1.25 µl |
| total | 50 µl |

4.2. Add 50µl of the ligation mix to each Hi-C tube (250 µl total).

4.3. Incubate at 20°C for 1 hours in ThermoMixer (800 rpm).

#### 5. Reverse crosslinking:

5.1. Add 5 µl of 20 mg/ml (500 U/ml) proteinase K.

5.2. For **CI** and **D** controls: add 45 µl PBS x1 and 1 µl 20 mg/ml (500 U/ml) proteinase K to each control

5.3. Incubate at 65°C for 1 hour in ThermoMixer (800 rpm).

#### 6. DNA Purification:

6.1. Transfer the Hi-C sample to 2ml PhaseLock tube. For CI and D no PhaseLock needed.

6.2. To each PhaseLock tube add 0.5ml saturated phenol:chloroform (1:1 pH=8.0).

6.3. Add the 0.25 ml sample to the PhaseLock tube.

6.4. To **CI** and **D** add 100 µl phenol:chloroform.

6.5. Vortex for 30-60 sec to obtain a homogenous milky solution.

6.6. Centrifuge at 16,000 x g for 5 min RT.

6.7. Transfer the aqueous phase with DNA to a new 1.7ml Eppendorf tube.

6.8. * Skip this stage if starting material is less than 50k cells: Transfer 5 µl from the sample to a new 1.7 ml Eppendorf tube for **Ligation control (L).**

**6.9. Ethanol precipitation:**

i. To each Hi-C sample tube (not to **CI**, **D**, **L**) add: 25 µl 3M sodium acetate pH=5.2 and 650 µl 100% ethanol, and vortex gently.

ii. Incubate at -80°C for 20 min.

iii. Centrifuge experiment tubes 16,000 x g for 20 min at 4°C.

iv. Dissolve the pellet in 100 µl of TLE.

v. Quantify DNA concentration using a fluorimeter dsDNA high sensitivity kit (DeNovix) on 1µl of sample.

6.10. While samples are precipitating, check CI, D and L controls on agarose gel:

i. Add DNA loading dye to 20 µl **CI** and **D**, and 5 µl **L**.

ii. Run samples on 1.2-1.5% agarose gel.

#### 7. Biotin removal:

7.1. Prepare biotin removal mix:

| **Biotin removal** | **Per sample** |
|---|---|
| Hi-C sample (up to 7 ug) | 100 µl |
| 10 x NEB buffer 2.1 | 12 µl |
| 10mM dATP | 0.25 µl |
| 10mM dGTP | 0.25 µl |
| 3000U/ml T4 DNA polymerase NEB | 6 µl |
| water | 1.5 µl |

7.2. Split each sample into two 60 µl samples to accommodate incubation in a PCR machine.

7.3. Incubate at 20°C for 1 hour, followed by 75°C for 20 minutes to inactivate T4 DNA polymerase.

7.4. Keep at 4°C overnight.

#### 8. DNA sonication:

8.1. Combine reactions from the same sample (total 120 µl) and transfer to a Covaris microtube for sonication.

8.2. Shear the DNA to 200-300bp using a Covaris sonicator at 20°C:

| **Covaris model** | **M220** |
|---|---|
| Peak Incident Power (Watt) | 50W |
| Duty Cycle/Duty factor | 20% |
| Intensity | ---- |
| Cycles per Burst | 200 |
| Treatment time (seconds) | 110 |
| Set Mode | |
| Pre-use degassing | --- |

#### 9. Biotin Pulldown with Streptavidin C1 beads:

* All work with beads is performed with **Low Binding** tubes and tips.

9.1. Vortex MyOne™ Streptavidin C1 beads and transfer 10 µl beads to a low-binding PCR tube. Do not add the DNA yet.

9.2. Wash the beads with 200 µl of Tween wash buffer (TWB) by pipetting up and down.

9.3. Incubate for 3 min at RT on a rocking platform.

9.4. Reclaim the beads against a magnetic rack for 1 minute and discard the supernatant.

9.5. Resuspend beads in 120µl of 2 x binding buffer (BBx2).

9.6. Add 120 µl of the sonicated Hi-C sample.

9.7. Incubate the sample for 15 minutes at RT on a rocking platform.

9.8. Gently spin down and reclaim the beads against a magnetic rack for 1 min.

9.9. Discard the supernatant and resuspend in 200 µl of BBx1 (1:1 TLE: BBx2).

9.10. Reclaim the beads against a magnetic rack for 1 min.

9.11. Discard the supernatant and wash the beads with 100 µl TLE.

9.12. Reclaim the beads against a magnetic rack for 1 min.

9.13. Discard the supernatant and resuspend the beads in 50 µl TLE.

9.14. Quantify the amount of DNA bound to streptavidin beads using a fluorimeter dsDNA ultra-high sensitivity kit (DeNovix) on 1 µl of sample.

#### 10. NEB end repair & A-tailing, and Illumina adaptor ligation

10.1. Create mix according to the NEBNext Ultra II protocol as follows:

| **End repair & A-tailing** | **Per sample** |
|---|---|
| Biotin pull down DNA | 50 µl |
| NEBNEXT Ultra II end prep reaction buffer | 7 µl |
| NEBNEXT Ultra II end prep enzyme mix | 3 µl |
| total | 60 µl |

10.2. Incubate reactions in the PCR at 20°C for 30 min to add adenine on the 3'end and then at 65°C for 30 min to inactivate enzymes with heated lid set to 75°C.

10.3. Proceed to adaptor ligation immediately. Add to each tube:

| Adaptor ligation | Per sample |
|---|---|
| End prep reaction mix | 60 µl |
| NEBNext Adaptor for Illumina** | 2.5 µl |
| NEBNext Ultra II Ligation master mix | 30 µl |
| NEBNext Ligation enhancer | 1 µl |
| total | 93.5 µl |

** Dilute the NEBNext adaptor for Illumina in TLE 1:10 if the DNA concentration measured by Denovix is 10-60 ng, 1:25 if 1-10 ng, or 1:40 if less than 1 ng material.

10.4. Mix well and incubate 20°C for 15 min in PCR with heated lid off.

10.5. Add 1 µl of USER Enzyme to the ligation mixture. Mix well and incubate at 37°C for 15 min with heated lid set to 47°C.

10.6. Reclaim the beads against a magnetic rack for 1 min. Discard the supernatant.

10.7. Resuspend the beads in 100 µl of BBx1.

10.8. Reclaim the beads against a magnetic rack for 1 min. Discard the supernatant.

10.9. Resuspend the beads in 100 µl of TLE.

**!!!**

For starting material >50k cells proceed to the next step. If your sample has less than 50k cells, skip to step 11.b.

**!!!**

10.10. Reclaim the beads against a magnetic rack for 1 min. Discard the supernatant, and resuspend beads in 15 µl TLE.

#### 11.a. Calibration PCR: (For samples with **at least 50k cells**)

11.a.1.Mix the following reagents for calibration PCR (pay attention to only use half the Hi-C sample at this stage):

| **Calibration PCR** | **Per sample** |
|---|---|
| KAPA HiFi HotStart ReadyMix | 12.5 µl |
| Adaptor ligated DNA fragments | 7.5 µl |
| NEBNext Multiplex Oligos for Illumina | 5 µl |
| Total | 25 µl |

11.a.2.Run the PCR program with pauses after different number of cycles:

| **STEP** | **TEMP** | **TIME** |
|---|---|---|
| 1 | 98°C | 60 seconds |
| 2 | 98°C | 15 seconds |
| 3 | 60°C | 30 seconds |
| 4 | 72°C | 40 sec |
| 5 | Go to 2 N times | ***X 15 – 21 times*** |
| 6 | 72°C | 3 minutes |
| 7 | 4°C | ∞ |

11.a.3.Quantify amount of DNA with fluorimeter dsDNA high sensitivity kit (DeNovix):

- If starting with 10-60ng DNA take 1 µl aliquot after 6,8,10,12 PCR cycles.
- If starting with 10-10ng DNA take 1 µl aliquot after 8,10,12,14 PCR cycles.
- If starting with less than 1ng DNA take 1 µl aliquot after 12,14,16,18 and 21 cycles

The Goal is to find the PCR cycle where DNA amount is >25ng/ µl.

11.a.4.Proceed to step 12 – ClaI digestion.

#### 11.b. (For samples with **less than 50k cells**)

11.b.1. Reclaim the beads against a magnetic rack for 1 min.

11.b.2. Discard the supernatant and resuspend beads in 7.5 µl TLE.

11.b.3. Prepare the sample for PCR amplification:

11.b.4.

| **Library-preparation PCR** | **Per sample** |
|---|---|
| KAPA HiFi HotStart ReadyMix | 12.5 µl |
| Adaptor ligated DNA fragments | 7.5 µl |
| NEBNext Multiplex Oligos for Illumina | 5 µl |
| Total | 25 µl |

11.b.5. Run the PCR program for 10 cycles:

| **STEP** | **TEMP** | **TIME** |
|---|---|---|
| 1 | 98°C | 60 seconds |
| 2 | 98°C | 15 seconds |
| 3 | 60°C | 30 seconds |
| 4 | 72°C | 40 sec |
| 5 | Go to 2 | ***X 9*** |
| 6 | 72°C | 3 minutes |
| 7 | 4°C | ∞ |

11.b.6. Quantify amount of DNA by measuring 1 µl with fluorimeter dsDNA high sensitivity kit (DeNovix).

a. If concentration is less than ~20n ng/ µl, return sample to PCR for 2 more cycles (12 cycles total).

b. If concentration is around ~20 ng/ µl and higher: Reclaim the beads on a magnetic rack for 1 minute, and transfer the supernatant containing the PCR products to a new 200 µl low-binding PCR tube. Store in 4°C until after ClaI control validation (step 12)

11.b.7. Resuspend the streptavidin beads in 100 µl of BBx1.

11.b.8. Reclaim the beads against a magnetic rack for 1 min. Discard the supernatant.

11.b.9. Resuspend the beads in 100 µl of TLE.

11.b.10. Reclaim the beads against a magnetic rack for 1 min. Discard the supernatant and resuspend beads in 7.5 µl TLE.

11.b.11. Prepare the sample for a second round of PCR amplification:

* Make sure to use the exact same indexes as the initial amplification.

| **Second PCR** | **Per sample** |
|---|---|
| KAPA HiFi HotStart ReadyMix | 12.5 µl |
| Adaptor ligated DNA fragments | 7.5 µl |
| NEBNext Multiplex Oligos for Illumina | 5 µl |
| Total | 25 µl |

11.b.12. Repeat the PCR program for 12 cycles, and digest with ClaI according to step 12 to validate successf µl ligation.

11.b.13. Continue with the PCR products from the first PCR (from stage 14 to AMpure size selection

#### 12. ClaI digestion:

12.1. Use the remaining calibration-PCR product to digest with ClaI:

| **ClaI digestion** | **Per sample** |
|---|---|
| Hi-C DNA sample | 21 µl |
| Cutsmart x 10 | 4 µl |
| water | To 40 µl |

12.2. Separate each sample to 2 PCR tubes each with 20 µl.

12.3. Add 0.5 µl ClaI to the (+) tube and 0.5 µl water to the (-) tube.

12.4. Incubate at 37°C for 30 minutes and run samples on 1.2-1.5% agarose gel.

12.5. Compare ClaI digested and undigested DNA - The undigested should be around 350bp and the digested should be smaller size a smear.

Proceed to production PCR only if DNA is digested by ClaI.

#### 13. PRODUCTION PCR and primer dimer removal:

13.1. Mix the following ingredients similar to calibration PCR:

| Ingredient | Volume |
|---|---|
| KAPA HiFi HotStart ReadyMix | 12.5 µl |
| Adaptor ligated DNA fragments | 7.5 µl |
| NEBNext Multiplex Oligos for Illumina | 5 µl |
| Total | 25 µl |

13.2. Set the PCR program number of cycles as determined by the calibration PCR to the lowest PCR cycle with DNA concentration > 25 ng/ µl:

| Step | Temp | Time |
|---|---|---|
| 1 | 98°C | 60 seconds |
| 2 | 98°C | 15 seconds |
| 3 | 60°C | 30 seconds |
| 4 | 72°C | 45 sec |
| 5 | Go to 2 N times | as determined in calibration PCR |
| 6 | 72°C | 3 minutes |
| 7 | 4°C | ∞ |

13.3. Reclaim the beads on a magnetic rack for 1 minute, and transfer the supernatant containing the PCR products to a new 200 µl low-binding PCR tube.

#### 14. Size fractionation using AMpure XP:

14.1. Purify the amplified Hi-C library from the supernatant using AMpure XP beads as follows:

i. Let AMpure XP mixture get to RT and mix thoroughly via vortex prior to using it.

ii. Add AMpure beads to the PCR product at a 1:1 ratio (25 µl:25 µl)

iii. Incubate at RT on a rocking platform for 5 min.

iv. Reclaim the beads on a magnetic rack for 1 minute, and discard supernatant.

v. Wash twice with 50 µl freshly made 80% ethanol.

vi. Air dry the beads for 5 minutes.

vii. Resuspend the beads in 20 µl TLE buffer to elute the DNA. Incubate RT on rocking platform 5 min.

viii. Reclaim the beads with a magnetic rack for 1 minute.

ix. Transfer the supernatant, containing the final Hi-C library, to a 1.5 ml Eppendorf tube.

14.2. Determine final Hi-C library concentration by measuring 1 µl with fluorimeter dsDNA high sensitivity kit (DeNovix), and determine the quality using TapeStation.

#### Buffers:

*all water should be mol. Biol. Grade

| **1X Lysis buffer** | **Stock** | **50 ml** |
|---|---|---|
| 10 mM Tris-HCl (pH=8.0) | 1 M (pH=8.0) | 500 µl |
| 10 mM NaCl | 5 M | 100 µl |
| 0.2% Igepal CA-630 (NP40) | 10% | 1000 µl |
| Water | to 50 ml | ~48.4 ml |
| --- Store at 4°C | | |

| **Tween Wash Buffer (TWB)** | **Stock** | **100 ml** |
|---|---|---|
| 5 mM Tris-HCl (pH=8.0) | 1M (pH=8.0) | 500 µl |
| 0.5 mM EDTA | 500 mM | 100 µl |
| 1 M NaCl | 5 M | 20 ml |
| 0.05% Tween | 100% | 50 µl |
| Water | to 100 ml | ~79.4 ml |

| **2X Binding Buffer (BB)** | **Stock** | **100 ml** |
|---|---|---|
| 10 mM Tris-HCl (pH=8.0) | 1 M (pH=8.0) | 1 ml |
| 1 mM EDTA | 500 mM | 200 µl |
| 2 M NaCl | 5 M | 40 ml |
| Water | to 100 ml | ~58.8 ml |

| **TLE (pH 8.0)** | **Stock** | **100 ml** |
|---|---|---|
| 10 mM Tris-HCl | 1 M | 1000 µl |
| 0.1 mM EDTA | 500 mM | 20 µl |
| Water | | ~99 ml |

#### Reagents and Equipment:

| Reagent | Company | Catalog number |
|---|---|---|
| Halt Protease Inhibitor Cocktail (100X) | Thermo | 78429 |
| Biotin-14-dATP | Jena bioscience | NU-835-BIO014-L |
| Proteinase K (Fungal) | NEB | P810S |
| DNA polymerase I, large (Klenow) fragment | NEB | M0210S |
| T4 DNA ligase 400U/ µl | NEB | M0202L |
| T4 DNA polymerase | NEB | M0203L |
| 10X ligation buffer | NEB | 46300-018 |
| T4 polynucleotide kinase | NEB | M0201 |
| Dynabeads® MyOne™ Streptavidin C1 | Invitrogen | 650.01 |
| Agencourt AMPure® XP | Beckman Coulter | A63881 |
| DNA LoBind tubes, 1.5 ml | Axygen | 81876MCT175LC |
| SUPERLADDER 2100 100BP DNA LADDER | Bio-Lab | 9597580SL2100 |
| 1 kb DNA marker | NEB | N3232S |
| Dynal MPC™-S Magnetic Particle Concentrator | Invitrogen | 120.20D |
| NEBNext Ultra II DNA Library | NEB | E7645S |
| KAPA HiFi HotStart ReadyMix | Roche | KK2601 |
| NEBNext Multiplex Oligos for Illumina | NEB | E6440S |
| NEBuffer 3.1 | NEB | B7203S |
| 37% formaldehyde(FA) | Sigma Aldrich | 252549 |
| dNTP set | NEB | N0446S |
| Triton X-100 10% solution | Sigma | 93443 |
| NP40 | Sigma | I8896 |
| Poly(ethylene glycol)4000 | Sigma | 95904 |
| Phase lock heavy gel | 5prime | 2302830 |
| Phenol – chloroform – isoamyl alcohol mixture | Bio-Lab | 1697234400 |
| (10%) SDS Solution | Biological Industry | 01-890-1B |
| DSDNA HIGH SENSITIVITY 1000 RXN-DSDNA-H2 | DeNovix | IM -DSDNA-H2 |
| DSDNA ULTRA HIGH SENSITIVI,1000 DSDNA-U2 | DeNovix | IM -DSDNA-U2 |
